# Discovery of a unique pathway for glutathione utilization in Francisella

**DOI:** 10.1101/2023.02.02.526638

**Authors:** Yaxi Wang, Hannah E. Ledvina, Catherine A. Tower, Stanimir Kambarev, Elizabeth Liu, James C. Charity, Lieselotte S.M. Kreuk, Qiwen Chen, Larry A. Gallagher, Matthew C. Radey, Guilhem F. Rerolle, Yaqiao Li, Kelsi M. Penewit, Serdar Turkarslan, Shawn J. Skerrett, Stephen J. Salipante, Nitin S. Baliga, Joshua J. Woodward, Simon L. Dove, S. Brook Peterson, Jean Celli, Joseph D. Mougous

## Abstract

Glutathione (GSH) is an abundant metabolite that can act as a signal, a nutrient source, or serve in a redox capacity for intracellular pathogens. For Francisella, GSH is thought to be a critical in vivo source of cysteine; however, the cellular pathways permitting GSH utilization by Francisella differ between strains and have remained poorly understood. Using genetic screening, we discovered a unique pathway for GSH utilization in Francisella spp. Whereas prior work suggested GSH catabolism initiates in the periplasm, the pathway we define consists of a major facilitator superfamily member that transports intact GSH and a previously unrecognized bacterial cytoplasmic enzyme that catalyzes the first step of GSH degradation. Interestingly, we find that the transporter gene for this pathway is pseudogenized in pathogenic Francisella, explaining phenotypic discrepancies in GSH utilization among Francisella spp. and revealing a critical role for GSH in the environmental niche of these bacteria.

## Introduction

It is increasingly appreciated that the success of bacterial pathogens relies on sophisticated strategies for scavenging nutrients from their hosts. These “nutritional virulence factors” can include mechanisms for manipulating the host to drive nutrient availability^1^. For example, some intracellular pathogens hijack autophagic or proteolytic cellular machinery to release amino acids that can be exploited as carbon and energy sources^2,3^. Other pathogens compete effectively with the host for nutrients that are available as a result of normal host physiology.

One metabolite present in particularly high abundance inside host cells is the tri-peptide glutathione (γ-L-glutamyl-L-cysteinyl-glycine, GSH). GSH and its oxidized counterpart GSSG play crucial roles in multiple essential processes including maintaining redox homeostasis, defense against reactive oxygen species, and protein iron-sulfur cluster synthesis ^4^. Perhaps as a result of the ubiquity and high concentration of GSH in the cytosol of eukaryotic cells, certain intracellular pathogens couple GSH sensing to virulence factor induction^5^. Notable examples include *Burkholderia pseudomallei*, which induces type VI secretion transcription following GSH sensing by the VirAG two-component system, and *Listeria monocytogenes*, which senses GSH through PrfA, leading to the activation of a set of critical virulence determinants^6,7^. Other pathogens, including *Hemophilus influenza* and *Streptococcus* spp. rely on co-opted host GSH to defend against oxidative stress^8,9^.

In contrast to these pathogens, Gram-negative proteobacteria belonging to the genus *Francisella* are sulfur amino acid auxotrophs and catabolize host GSH as a source of organic sulfur. A transposon screen of *Francisella tularensis* subspecies *holarctica* LVS (*F. tularensis* LVS) revealed that the periplasmic enzyme γ-glutamyl transpeptidase (GGT), which cleaves GSH into glutamate and cysteine–glycine (Cys–Gly), is essential for intracellular replication of this organism^10^. Using a similar approach, our laboratory identified an inner membrane proton-dependent oligopeptide transporter-family (POT) protein that imports Cys–Gly^11^. We named this protein DptA and demonstrated that, consistent with its critical role in GSH catabolism, *F. tularensis* LVS Δ*dptA* is defective in intracellular replication. However, we found that inactivation of *ggt* or *dptA* neither attenuates intracellular growth nor compromises GSH catabolism in a closely related *Francisella* strain, *F. tularensis* subsp. *novicida* (*F. novicida*). Rather, we identified a predicted γ-glutamylcyclotransferase enzyme in *F. novicida*, ChaC, that participates in GSH catabolism and is required for robust *F. novicida* growth in media containing GSH as the sole organic sulfur source (GSH media).

Despite these additions to our understanding of GSH metabolism in *Francisella*, two lines of evidence suggested that it remained incomplete. First, if GSH breakdown by Ggt and ChaC represented the only entry points into GSH catabolic pathways in *F. novicida*, a strain lacking these enzymes should be unable to grow in media containing GSH as a sole cysteine source. On the contrary, we found that *F. novicida* Δ*ggt* Δ*chaC* grows in such media, albeit not at wild-type levels^11^. Second, although *ggt, dptA* and *chaC* are present and expected to be functional in both *F. novicida* and *F. tularensis* LVS, inactivation of *ggt* only produces a growth defect in GSH media in *F. tularensis* LVS. Together, these observations led us to hypothesize that additional pathways for GSH catabolism remain to be uncovered in *Francisella*.

In this study, we employed Tn-seq to identify Ggt-independent pathways important for GSH utilization in *Francisella*. Through this analysis, we discovered that *F. novicida* possesses a previously unrecognized pathway for GSH utilization that consists of an outer membrane porin, an inner membrane transporter of intact GSH belonging to the major facilitator superfamily and a cytoplasmic glutamine amidotransferase family enzyme capable of initiating degradation of the molecule. We show that this pathway is mutationally inactivated in pathogenic *Francisella* spp., but widely conserved in members of the genus that are believed to inhabit an environmental niche. Our work thus has implications for the evolution of pathogenesis within *Francisella*, and provides evidence that the natural lifecycle of non-pathogenic *Francisella* likely includes replication within a GSH-rich habitat, such as the cytosol of unicellular eukaryotes.

## RESULTS

### Tn-Seq reveals genes required for GSH utilization in *F. novicida* U112

Although Ggt is required for the growth of *F. tularensis* LVS in GSH medium, we previously found that its inactivation does not similarly impede *F. novicida* growth on this substrate^11^. Moreover, using the genome of *F. novicida*, we were unable to identify additional characterized GSH catabolism pathways that are absent from *F. tularensis* LVS. This conundrum motivated us to undertake an unbiased approach for discovering GSH catabolism pathways in *F. novicida*. To this end, we generated transposon mutant libraries of *F. novicida* in the wild-type and Δ*ggt* backgrounds, and used transposon mutant sequencing to compare gene insertion frequencies for each library grown in media containing GSH versus cysteine as the sole sulfur source (Figures 1A-1C). Our decision to employ Δ*ggt* rather than Δ*chaC* in this experiment was motivated by our recent observation that, while *ggt* is not required for *F. novicida* growth when GSH is in excess (100 μM), growth of the Δ*ggt* strain is slightly retarded when the concentration of GSH limits growth (Figure S1). Strains lacking Δ*chaC* grow to wild-type levels under both conditions, suggesting that Ggt has a larger role in GSH catabolism in our *in vitro* culturing conditions.

**Figure 1.**
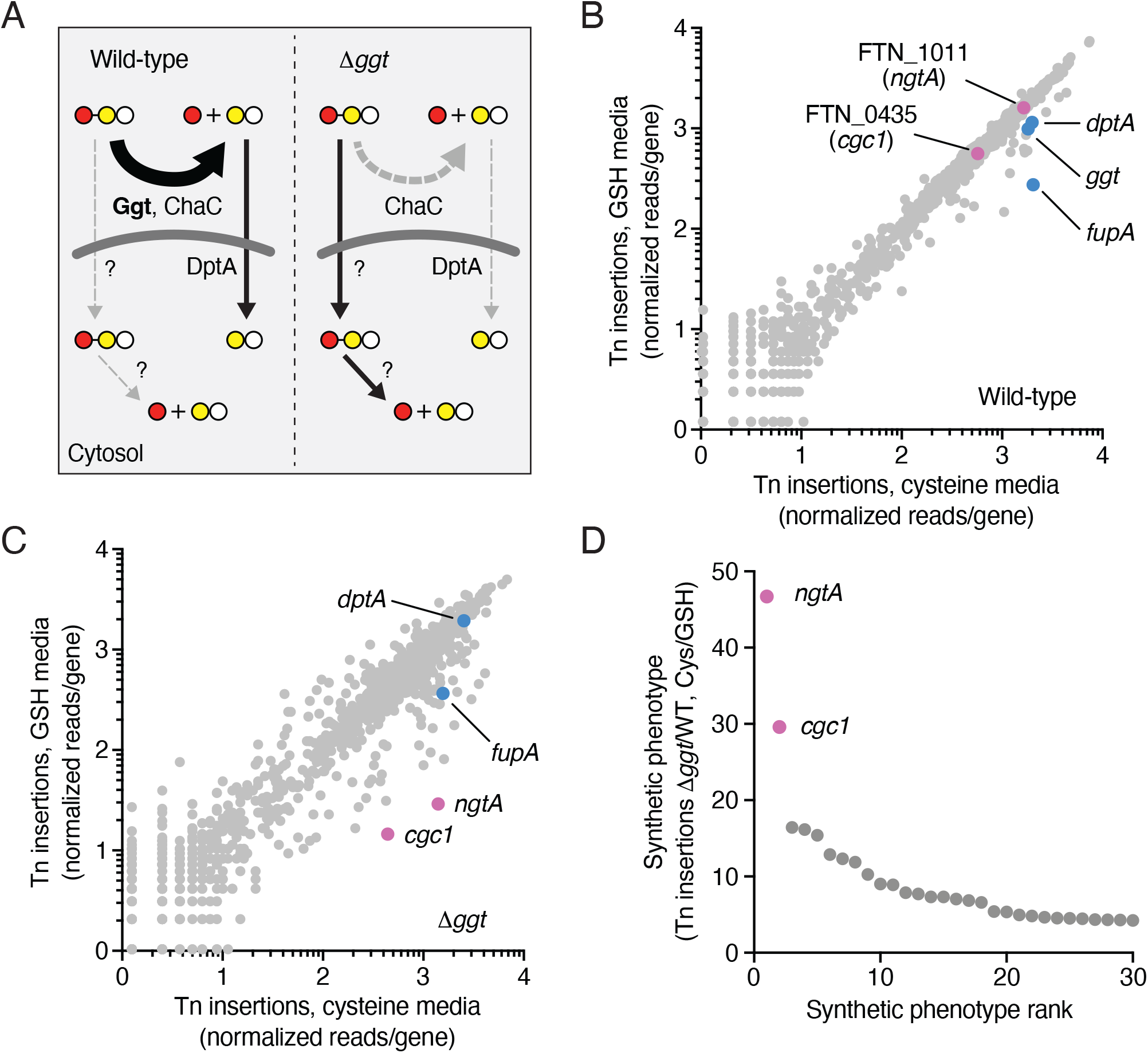
Tn-seq for discovery of *F. novicida* genes with synthetic phenotypes during growth on GSH. (A) Schematic illustrating known and unidentified potential GSH catabolism pathways and their products (Glu, red circle; Cys-Gly, yellow and white circles) in the two genetic backgrounds employed in our screen. The heavy arrow depicted for the wild-type background emphasizes the primary conversion pathway (left, Ggt-mediated) while the dashed arrow indicates the residual GSH cleavage mediated by ChaC in the absence of Ggt (right). (B,C) Results of Tn-seq screen to identify genes required for growth of *F. novicida* on GSH medium in the wild-type (B) and Δ*ggt* backgrounds (C). Genes with the greatest difference in transposon insertion reads between growth in GSH and cysteine media in the Δ*ggt* background (purple) and other genes shown previously or in this study to participate in GSH uptake or catabolism (blue) are indicated. (D) Rank order depiction of the strength of the synthetic phenotype for genes important for growth of *F. novicida* Δ*ggt* in GSH medium. Rank order was calculated by dividing the ratio of transposon insertion frequency obtained for each gene during growth on GSH compared to growth on cysteine using the *F. novicida* Δ*ggt* background by the same ratio obtained using the wild type background.

Our Tn-seq screen led to the identification of many genes important for the growth of *F. novicida* Δ*ggt* specifically in GSH media. Among the 40 top hits in the Δ*ggt* background – corresponding to a three-fold insertion frequency difference between cysteine and GSH as the sole sulfur source – only five were shared with wild-type (Tables S1 and S2). Interestingly, while important for growth in GSH media specifically in the Δ*ggt* background, *chaC* was not among the strongest hits we observed (Table S2), supporting our earlier observation that Δ*ggt* Δ*chaC* can propagate in GSH media. Also consistent with our prior findings, in the wild-type strain, the fitness cost of inactivating *ggt* or *dptA* in GSH media was modest (Figure 1B).

To highlight Ggt-independent pathways for GSH catabolism, we ranked *F. novicida* genes by the strength of their synthetic (Δ*ggt* versus wild-type) growth phenotype in GSH media. Two genes ranked substantially higher in this analysis than other hits from our screen: FTN_1011 and FTN_0435 (Figure 1D). These two genes were also those with the greatest difference in insertion frequency between growth on GSH and cysteine as sole sulfur sources in the Δ*ggt* background (46.7- and 29.6-fold difference in normalized read counts, respectively) (Figure 1C and Table S2). Neither of these genes have been characterized, nor have functions been ascribed to any close homologs. Thus, we hypothesized they could contribute to GSH catabolism through previously unknown mechanisms.

### Identification and characterization of GSH transporter NgtA

The strongest synthetic phenotype during growth in GSH medium belonged to open reading frame FTN_1011 – herein named *ngtA* (<u>n</u>ovicida <u>g</u>lutathione transporter <u>A</u>). NgtA is a member of the major facilitator superfamily (MFS) of transporters, and as is typical of these proteins, its predicted structure displays 12 transmembrane helices organized into two six-helix bundles connected by a flexible linker^12^. Within the MFS, NgtA was previously classified into the Pht family^13^. Interestingly, Pht family members are found exclusively in intracellular pathogens; in *Legionella pneumophila* and *Fransicella*, proteins in the family are important for intracellular replication by virtue of their role in amino acid transport or nucleoside transport ^13-17^. However, the sequence of NgtA is substantially divergent from characterized Pht family members (26% sequence identity shared with PhtA, the most closely-related characterized Pht family member), and its function and substrate are unknown.

We hypothesized that NgtA could be a transporter of GSH. To test this hypothesis, we first generated in-frame deletion mutants of *ngtA* in the *F. novicida* wild-type and Δ*ggt* backgrounds. As predicted by our Tn-seq results, inactivation of *ngtA* in the wild-type background did not affect *F. novicida* growth in GSH medium (Figure 2A). However, growth of *F. novicida* Δ*ggt* Δ*ngtA* was strongly impaired in GSH medium. This growth defect could be complemented by repairing the deletion of *ngtA* via allelic exchange, and inactivation of *ngtA* did not cause growth defects in media containing cysteine as a sole sulfur source in either background (Figure S2A). To determine whether NgtA contributes to GSH uptake in *F. novicida*, we mixed cysteine-starved strains with ^3^H-GSH ([Glycine-2-^3^H]-GSH) and measured cell-associated radiolabel following a short incubation (Figure 2B). In the wild-type strain, NgtA inactivation had no impact on ^3^H-GSH transport. Since Ggt and ChaC generate periplasmic Cys– Gly, which when transported into the cytoplasm by DptA would mask the potential role of NgtA in intact GSH transport, we next employed the Δ*ggt* Δ*chaC* background in these assays. As expected, these mutations diminished GSH transport; however, uptake of the labeled substrate dropped to levels approaching the limit of detection in *F. novicida* Δ*ggt* Δ*chaC* Δ*ngtA* (Figure S2B). This result supports the hypothesis that in the absence of GSH cleavage in the periplasm, the intact tripeptide can be transported to the cytoplasm via NgtA.

**Figure 2.**
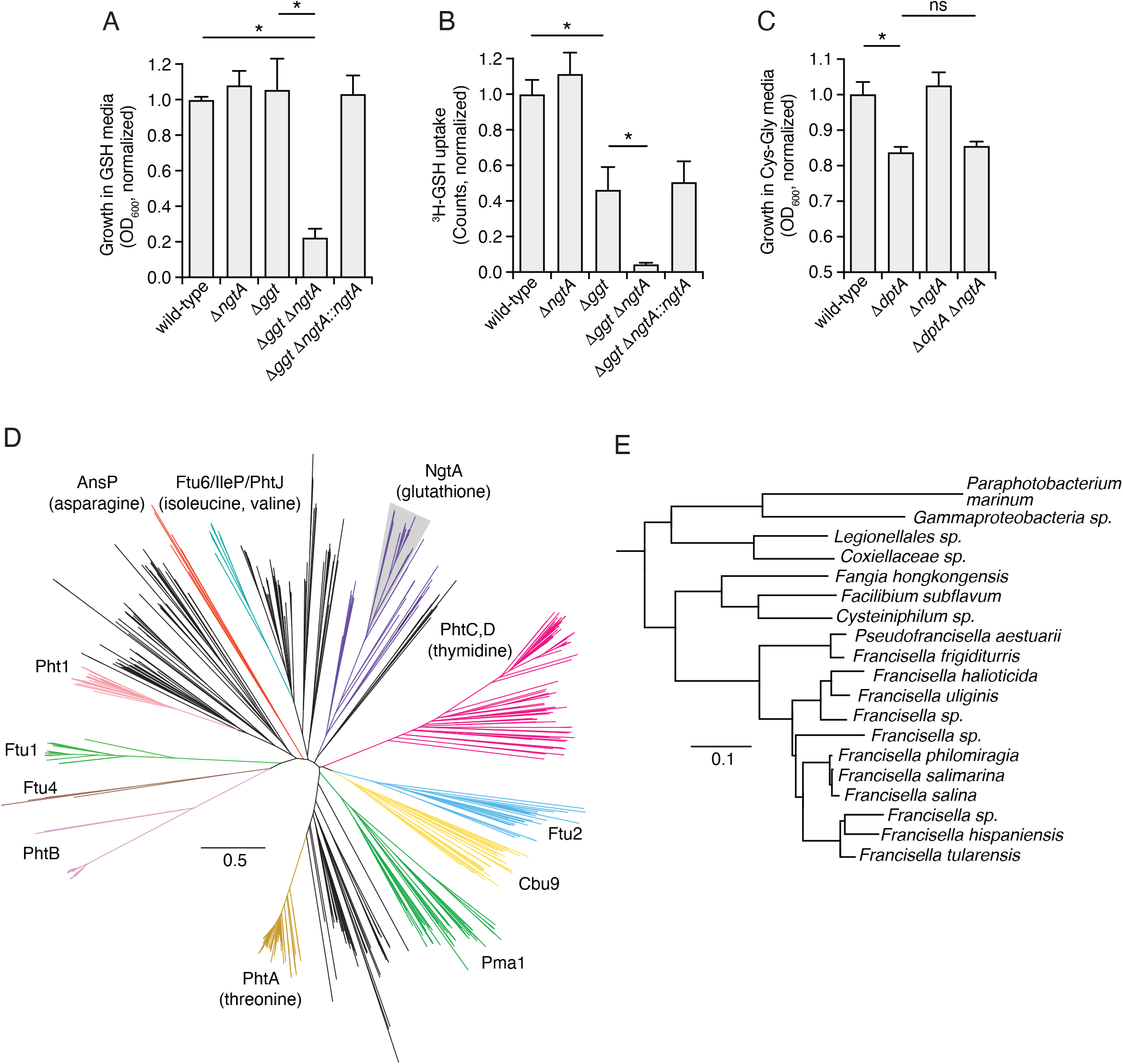
NgtA is a major facilitator superfamily protein that transports intact GSH in a Ggt-independent manner. (A) Normalized growth yield in GSH medium of the indicated *F. novicida* strains. (B) Quantification of the level of [Glycine-2-3H]-Glutathione (^3^H-GSH) uptake in the indicated strains of *F. novicida* after 45 min incubation. (C) Normalized growth yield of the indicated *F. novicida* strains after 36 hrs in defined medium containing Cys–Gly as a sole source of cysteine. (D) Neighbor-joining phylogeny of proteins from the Pht family of MFS transporters. Colored clades contain sequences identified in the original description of the family or subsequently characterized^13^. Representative proteins from the Chen *et al*. study or other reports are indicated by their respective clades, and transport substrate are indicated in parentheses when known. Candidate NgtA homologs are shown in purple, and the region of the phylogeny amplified in (E) is indicated (shading). (E) Neighbor-joining phylogeny of NgtA homologs in *Francisella* and related genera. Species names indicate the source of the protein sequences. Data in (A-C) represent mean ± s.d. Asterisks indicate statistically significant differences (Unpaired two-tailed student’s t-test.; *p<0.05, ns, not significant).

Our data left open the formal possibility that *F. novicida* possesses a third mechanism to generate Cys–Gly from GSH, and that the dipeptide is the transport substrate of NgtA. Notably, our laboratory previously reported the *F. novicida* Cys–Gly transporter DptA^11^. This strain exhibits only a partial growth defect in media containing Cys–Gly as a sole organic sulfur source (Cys–Gly media), suggesting that, indeed, other enzymes could support Cys–Gly transport (Figure 2C). However, inactivation of NgtA had no impact on *F. novicida* growth in Cys–Gly media in either the wild-type or Δ*dptA* backgrounds. Together, these data suggest that NgtA is a transporter with specificity for intact GSH.

In the initial report of the Pht family of MFS transporters, the NgtA homolog of *F. tularensis* was the only member of its cluster^13^. With many more genome sequences now available, we asked whether homologs of this protein could be found in other species. Using PSI-BLAST with the sequence of NgtA from *F. novicida* as the seed, we collected all publicly available sequences encoding MFS proteins from the Pht family and constructed a phylogeny. We found that while many of the clades in the phylogeny are dominated by sequences deriving from *Legionella* and *Coxiella* spp., the clade containing NgtA consists largely of sequences deriving from *Francisella* spp. and related *Thiotrichales* (Figures 2D and 2E). Our inability to identify NgtA orthologs more broadly suggests that this mechanism of transporting GSH may be an adaptation particularly exploited by organisms in this group.

### A cytoplasmic glutamine amidotransferase family enzyme that initiates GSH degradation

The finding that a GSH transporter can facilitate *F. novicida* growth in GSH media in the absence of Ggt and ChaC implies that this organism must encode cytoplasmic proteins capable of initiating GSH catabolism. The gene with the second strongest synthetic phenotype in our transposon mutant screen, FTN_0435, encodes a predicted glutamine amidotransferase (GATase). Most GATase proteins function in biosynthetic reactions in which the amido group from glutamine is transferred to an acceptor substrate, generating glutamate and an aminated product^18^. However, a limited number of GATase domain-containing proteins instead function as catabolic enzymes that cleave γ-glutamyl bonds in assorted substrates, releasing glutamate. These include enzymes that hydrolyze such substrates as the folate storage and retention molecule folylpoly-γ-glutamate, the spermidine degradation intermediate γ-glutamine-γ-aminobutyrate, GSH conjugates involved in glucosinolate synthesis, and notably, GSH itself^19-23^. The latter was found to occur in yeast and is catalyzed by the enzyme Dug3p^23^.

Structure modeling revealed that FTN_0435, herein named Cgc1 (<u>c</u>ytosolic <u>g</u>lutathione <u>c</u>atabolizing 1), shares an overall fold and a conserved predicted catalytic triad (C97, H184, E186) with class I GATases^24^ (Figure 3A). This is in contrast to the GSH-targeting enzyme of yeast, a class II GATase^23^. Nevertheless, we found that, as predicted by our Tn-seq results, Cgc1 is required for *F. novicida* Δ*ggt* growth in GSH medium (Figure 3B). Substitution of the predicted catalytic cysteine with alanine (*cgc1*^*C97A*^) recapitulated the phenotype of a *cgc1* deletion, supporting an enzymatic role for this protein in GSH catabolism. We thus asked whether Cgc1 encodes a cytoplasmic enzyme able to initiate GSH degradation.

**Figure 3.**
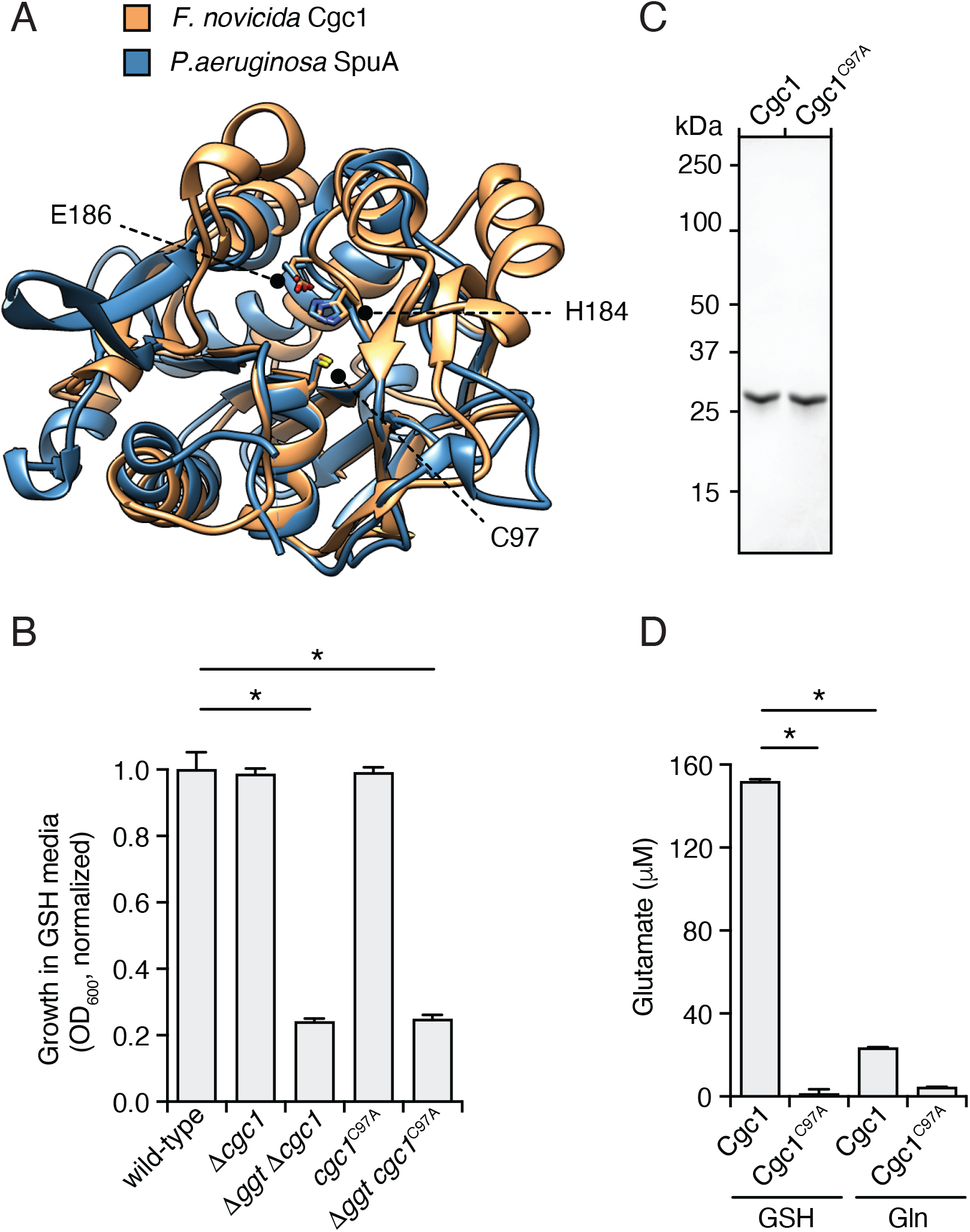
Cgc1 is a cytoplasmic glutamine amidotransferase (GATase) family protein that degrades GSH. (A) Alignment of the predicted structure of Cgc1(orange) and that of a characterized class I GATases, *P. aeruginosa* SpuA (blue, PDB: 7D4R). The conserved catalytic triad is indicated (numbers correspond to amino acid positions in *F. novicida* Cgc1). (B) Normalized 36 hrs growth yields of the indicated strains of *F. novicida*. (C) Coomassie stained SDS-PAGE analysis of purified Cgc1 and Cgc1^C97A^. (D) Glutamate released following 60 min incubation of purified Cgc1 or Cgc1^C97A^ (1 μM protein) with GSH or Gln (10 mM substrate). Data in (B) and (D) represent means ± s.d. Asterisks represent statistically significant differences (Unpaired two-tailed student’s t-test.; *p<0.05, ns, not significant).

To determine the substrate specificity of Cgc1, we used established *in vitro* assays to measure the activity of Cgc1 and Cgc1^C97A^ purified from *E. coli*. Biosynthetic GATase proteins exhibit glutaminase activity, generating glutamate and ammonia in the absence of their respective amido group-accepting substrates. However, we detected only a low level of glutamate accumulation following incubation of Cgc1 with glutamine. On the contrary, we readily detected glutamate released from GSH by Cgc1, and this product was not detected above background levels in reactions with Cgc1^C97A^ (Figures 3C and 3D). In total, these data support the hypothesis Cgc1 is a GATase that acts downstream of NgtA to initiate the degradation of GSH via cleavage into Glu and Cys–Gly.

Although our genetic and biochemical data strongly suggest that GSH is a physiological substrate of Cgc1, we noted the rate of glutamate release from the purified enzyme is low. In yeast, the GATase enzyme Dug3p acts in concert with two other proteins, Dug1p and Dug2p ^23^. Purified Dug3p is inactive *in vitro* unless bound to Dug2p, which allosterically activates the enzyme. Dug1p is a Cys–Gly specific peptidase that does not physically associate with the Dug2p–Dug3p complex. Cgc1 is encoded by the third gene in a predicted five gene operon. Examination of our Tn-seq results suggested that the genes encoded upstream of *cgc1* within this operon may also be important for growth on GSH in the *F. novicida* Δ*ggt* background (Figures S3A-C, Tables S1 and S2); however, we also considered that insertions within these genes may lead to polar effects on *cgc1*. To distinguish between these possibilities, we generated a conservative in-frame deletion in the first gene in the operon in *F. novicida* Δ*ggt* and measured the growth of this strain relative to *F. novicida* Δ*ggt* Δ*ngtA* in GSH media. This strain exhibited robust growth in GSH media (Figure S3D), strongly suggesting that polar effects underlie the apparent depletion of genes upstream of *cgc1* in our Tn-seq study, and moreover that Cgc1 does not require adjacently encoded proteins for its activity.

### Parallel pathways for GSH catabolism contribute to *F. novicida* intramacrophage growth

Previous studies indicate that *ggt* mutants of *F. tularensis* SCHU S4 and LVS are attenuated in virulence ^10,11,25-27^. This has led to the consensus in the field that GSH serves as an important source of organic sulfur for these bacteria during infection^1,5,28^. To our knowledge, the role of host GSH catabolism during *F. novicida* infection has not been examined. Unlike *F. tularensis* SCHU S4 and LVS, our results suggest that *F. novicida* may be capable of utilizing multiple pathways for GSH scavenging *in vivo*. To explore this possibility, we measured the growth of *F. novicida* strains lacking the function of one or both GSH uptake pathways in bone marrow-derived murine macrophages (BMMs). Interestingly, we found that only *F. novicida* strains in which both pathways are inactivated display a detectable intracellular growth defect (Figure 4A). We next examined the importance of the two GSH catabolism pathways in a more complex model of infection, a murine intranasal model^29^. At 48 hrs post-infection in the intranasal model, we observed a modest decrease in recovery of *F. novicida* Δ*ggt* from lung samples. However, in contrast to our macrophage infection study, no further decrease was detected when Δ*ggt* was combined with Δ*ngtA* or Δ*cgc1* (Figure 4B).

**Figure 4.**
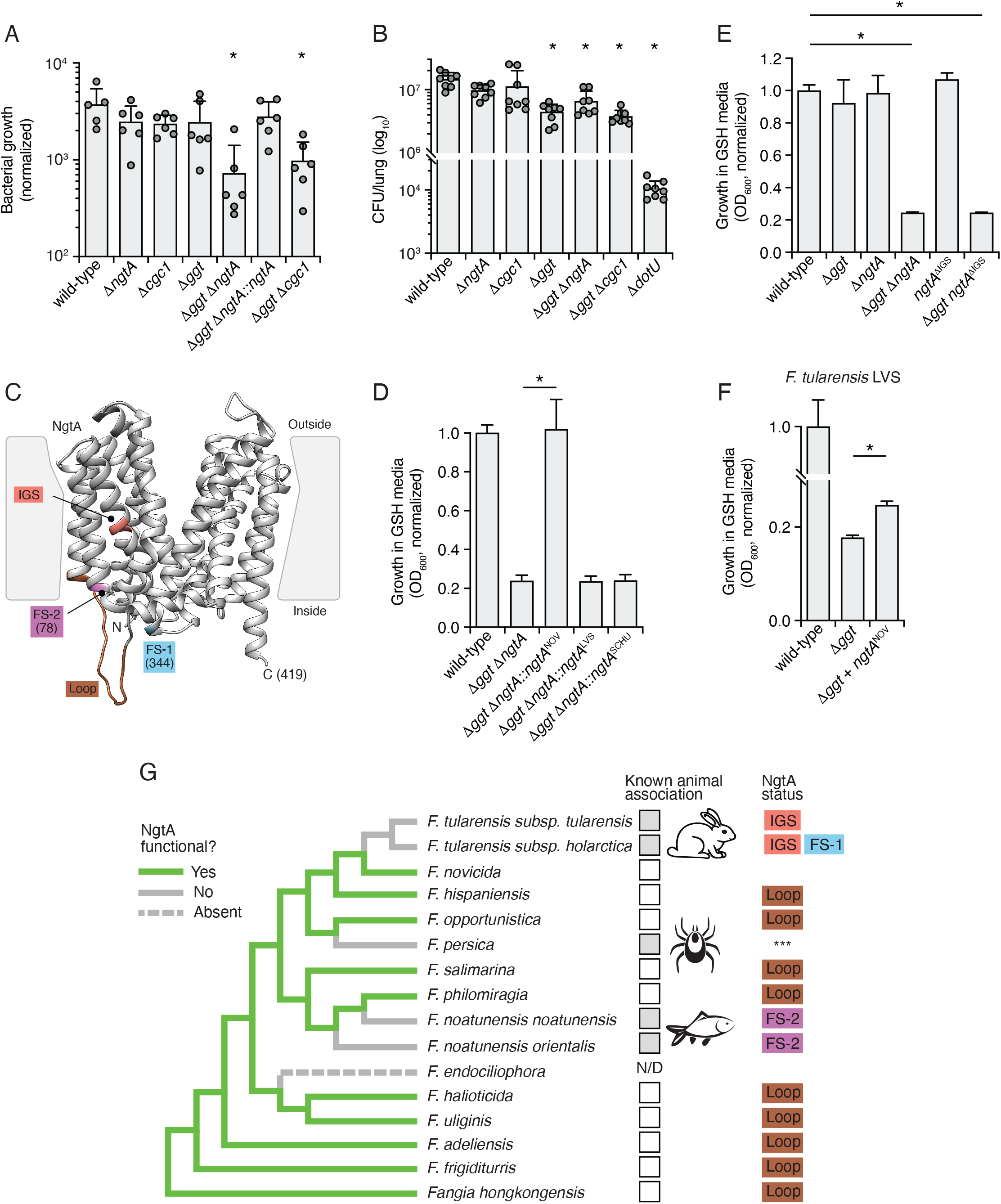
NgtA contributes to intramacrophage replication of *F. novicida* but is mutationally inactivated in pathogenic *Francisella* strains. (A) Normalized intracellular growth of the indicated strains of *F. novicida* in bone marrow-derived murine macrophages (24 hrs post-infection. (B) Bacterial burden in mouse lungs at 48 hrs post intranasal infection with ∼100 CFU of the indicated strains of *F. novicida*. (C) Structural model of *F. novicida* NgtA highlighting differences with NgtA in other *Francisella* species. The terminal residues resulting from truncating frameshift mutations (FS) indicated in parentheses. FS-1, location of truncation resulting from frameshift in *F. tularensis* subsp. *holarctica;* FS-2, location of truncation resulting from frameshift in *F. noatunensis*; loop, poorly conserved region with many differences between species; IGS, three-residue deletion found in *F. tularensis* subsp. *tularensis*. (D-F) Normalized growth yields in GSH medium of the indicated strains of *F. novicida* (D,E) or *F. tularensis LVS* (F). (G) Schematized phylogeny of *Francisella* species indicating predicted functionality of NgtA (functional, green; inactivated, solid grey; absent, dashed grey), animal association when known (mammal association indicated by rabbit schematic) and mutations present in *ngtA* (colors correspond to panel C). The *ngtA* sequence of *P. persica* contains many mutations (***) and *ngtA* appears to have been lost completely from *F. endociliophora*. Phylogenetic relationships derived from Vallesi *et al*.^42^. Data shown in (A) (B) and (D-F) represent as means ± s.d. Data points in (A-B) indicate technical replicates from 3 (A) or 4 (B) biological replicates conducted. Asterisks indicate statistically significant differences (A and B, one-way ANOVA followed by Dunnett’s; D-F, unpaired two-tailed student’s t-test; *p<0.05, ns, not significant.)

The results of our murine infection model study suggest that in the context of an animal infection, Ggt-mediated cleavage of GSH may be the primary mechanism by which *F. novicida* acquires organic sulfur. This is consistent with the observation that *ggt* mutants of *F. tularensis* SCHU S4 and *F. tularensis* LVS have significant virulence defects; however, it remained unclear why *ggt* inactivation alone is sufficient to inhibit *in vitro* growth in GSH medium in these other subspecies, but not in *F. novicida*. Furthermore, the magnitude of virulence defect for the Δ*ggt* background of *F. novicida* is qualitatively lower than that of *F. tularensis* SCHU S4 and LVS^10,11,25-27^. While investigating explanations for this difference, we found that the *ngtA* genes of SCHU S4 (*ngtA*^SCHU^), LVS (*ngtA*^LVS^), as well as those of other *F. tularensis* subsp. *Tularensis* and subsp. *holarctica* strains encode proteins with a three amino acid deletion relative to NgtA^NOV^. In the predicted structure of NgtA^NOV^, these amino acids (I113-G114-S115) reside within a transmembrane helix located in the core of the protein (IGS, Figure 4C). *F. holarctica ngtA* genes further contain a small 5′ in-frame deletion and a premature stop codon that removes the last two predicted transmembrane helices (FS-1, Figure 4C). Taken together with our current findings, these observations led us to hypothesize that NgtA, and thus GSH transport, is compromised in these pathogenic strains of *Francisella*. Indeed, we found that *F. novicida* Δ*ggt* carrying *ngtA*^SCHU^ or *ngtA*^LVS^ in place of *ngtA*^NOV^ demonstrated growth behavior matching *F. novicida* Δ*ggt ΔngtA* in GSH medium (Figure 4D). Furthermore, *F. novicida* Δ*ggt* carrying *ngtA*^*NOV*^ engineered to contain only the three-residue deletion found in *ngtA* alleles from human pathogenic strains was similarly unable to grow in GSH medium (Figure 4E). We also performed the converse experiment in *F. tularensis* LVS by over-expressing *ngtA*^NOV^ in the Δ*ggt* background. The expression of *ngtA*^NOV^ resulted in a small, but reproducible restoration of growth in GSH medium (Figure 4F). We speculate that the limited degree to which NgtA^NOV^ expression restores *F. tularensis* LVS GSH autotrophy could be the result of pseudogenization or regulatory alteration of elements downstream of NgtA that are important for efficient GSH catabolism.

Our finding that *ngtA* is inactivated in multiple *F. tularensis* subspecies prompted us to examine the nature and prevalence of mutations in *ngtA* amongst *Francisella* spp. more broadly. Interestingly, we found evidence supporting pseudogenization of *ngtA* in two additional lineages of animal-associated *Francisella*: the tick endosymbiont *F. persica* and the fish pathogens *F. noatunensis* and *F. orientalis* (Figures 4C and 4G). On the contrary, the *ngtA* sequences of *Francisella* without a known animal association bore mutations primarily restricted to a hypervariable cytoplasmic loop that are not expected to inactivate the transporter (Figures 4C and 4G). Given that we find evidence for repeated *ngtA* pseudogenization events limited to *Francisella* lineages adapted to animal hosts, our results suggest that intact GSH uptake is most beneficial for bacteria in this genus in the environment, perhaps during replication within unicellular eukaryotes.

### FupA is a porin that mediates GSH uptake

We were surprised to find *fupA* as a gene with highly differential transposon insertion frequency during growth on GSH versus cysteine as sole sulfur sources in both the wild-type and Δ*ggt* backgrounds of *F. novicida* (Figures 1B, 1C, Tables S1 and S2). FupA is a member of a family of paralogous predicted outer membrane proteins unique to *Fransicella* species, several of which, including FupA, are widely thought to mediate high affinity uptake of ferrous iron^30-32^. Despite this dogma, *F. tularensis* SCHU S4 Δ*fupA* exhibits a general growth defect in minimal media regardless of iron source or type, and proteoliposome assays using purified FupA provide evidence that it promotes membrane permeability^30,33^. We thus hypothesized that FupA may contribute to *F. novicida* growth in GSH media by facilitating GSH passage through the outer membrane.

To directly examine the role of FupA in GSH catabolism, we generated an in-frame deletion of *fupA* in the wild-type and Δ*ggt* backgrounds of *F. novicida*. Consistent with our Tn-seq results, deletion of *fupA* in both backgrounds resulted in a strong growth defect specifically in GSH media (Figure 5A). We then evaluated the role of FupA in GSH transport by measuring the impact of Δ*fupA* on cellular uptake of ^3^H-GSH by *F. novicida*. We found that in the absence of FupA, ^3^H-GSH uptake was reduced below levels observed in *F. novicida* Δ*ggt* (Figures 2B and 5B). Furthermore, inactivation of *ggt* in the Δ*fupA* background did not further reduce ^3^H-GSH uptake. These data support our hypothesis and further suggested that FupA could act as a general porin of *F. novicida*. Indeed, a prior analysis of predicted β-barrel proteins in *Francisella* did not identify clear homologs of previously characterized general porins^34^. In addition to serving as a conduit for the uptake of nutrients, a common feature of porins is that they present a vulnerability by providing entry to harmful molecules such as antibiotics and hydrogen peroxide ^35 36^. We found that *F. novicida* Δ*fupA* is significantly more resistant to hydrogen peroxide than the wild-type, further supporting its functional assignment as a porin of *F. novicida* (Figure 5C). These findings show that GSH accesses the periplasm of *F. novicida* via FupA, thus providing an explanation for the insertion frequency in *fupA* observed in our screen. In total, our genetic, biochemical and phenotypic data allow us to assemble a new, complete model for GSH transport and catabolism in *Francisella* (Figure 6).

**Figure 5.**
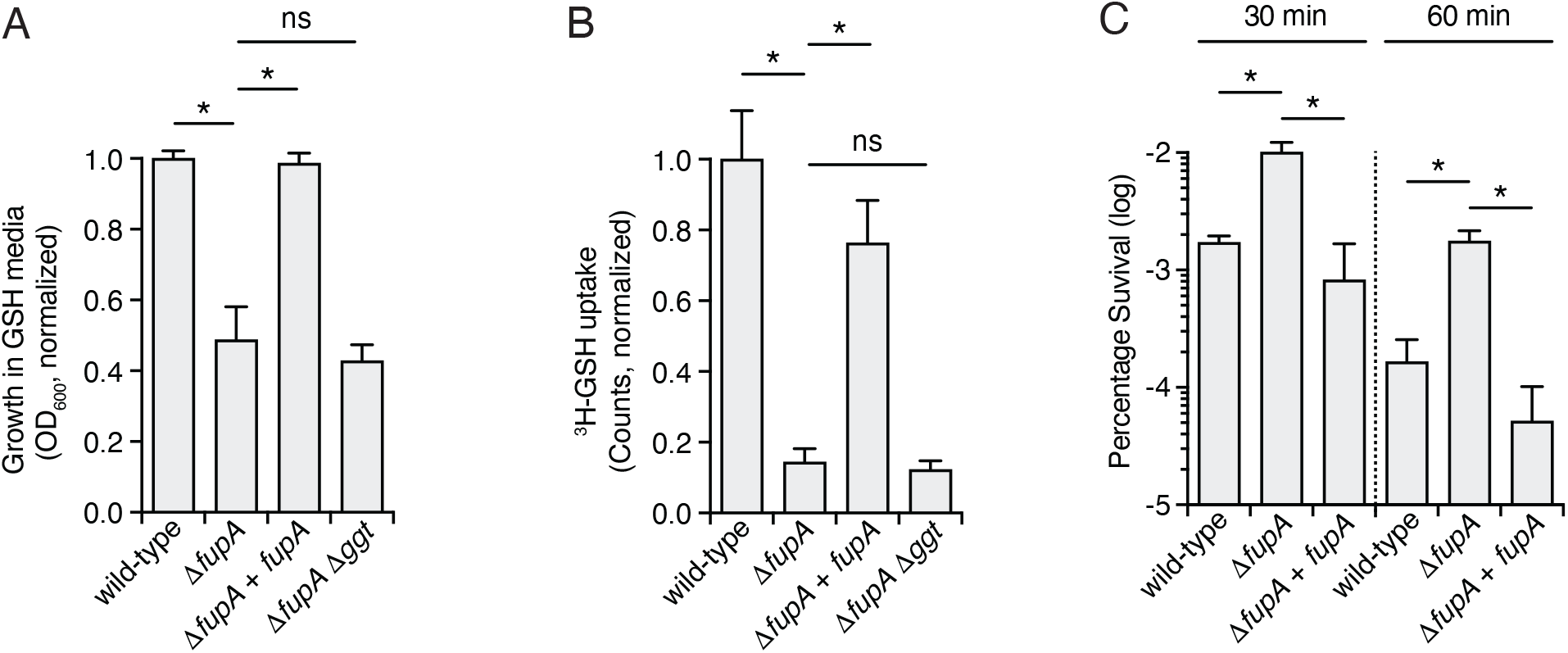
FupA is a porin required for GSH uptake in *F. novicida*. (A) Normalized growth in GSH mediume of the indicated strains of *F. novicida*. (B) Quantification of the level of ^3^H-GSH uptake in the indicated strains of *F. novicida* after 45 min incubation. (C) Survival of the indicated strains of *F. novicida* after incubation of mid-log phase cultures with 1.5 mM H_2_O_2_ for 30 min or 60 min. Data in (A-C) represent mean ± s.d. Asterisks indicate statistically significant differences (Unpaired two-tailed student’s t-test.; *p<0.05, ns, not significant).

**Figure 6.**
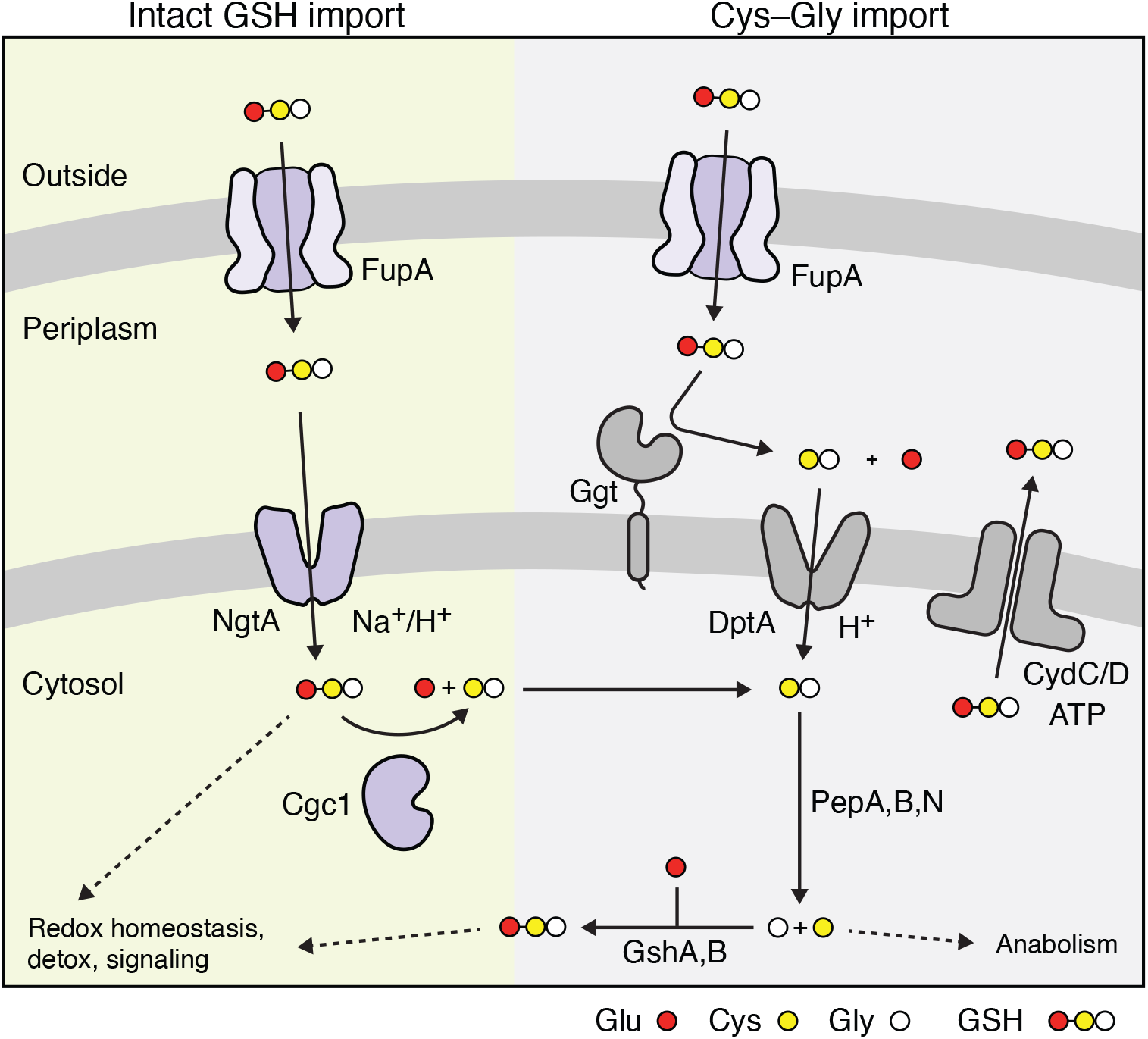
Comprehensive model of GSH transport and catabolism in *Francisella*. Both the pathways for import and cytosolic catabolism of GSH discovered in this study (left, green shading) and for periplasmic degradation of GSH and subsequent fate of imported Cys-Gly (right, grey shading) are indicated.

## Discussion

In this study, we report the finding that *Francisella* spp. encode a previously uncharacterized, Ggt-independent pathway for GSH uptake that has been lost in each established animal-colonizing lineage of the genus. On one hand, the correlation between inactivation of the GSH transporter encoding gene *ngtA* and adaptation to animal association is counterintuitive, as GSH is only present at sufficient concentrations to be useful as a source of sulfur in host-associated environments. However, we found evidence that in *F. novicida*, a species without a known physiological animal host, NgtA and the intracellular GSH-degrading enzyme Cgc1 work in concert with Ggt to support intramacrophage replication. This was not observed in a mouse model of infection, where a range of cell types are infected^37^. Macrophages share many features in common with amoeba, an abundant and widely distributed group of unicellular eukaryotes^38-40^. While the environmental niches colonized by non-pathogenic *Francisella* species remain largely uncharacterized, several species have been isolated from single-celled eukaryotes, including the deeply branching species *F. adeliensis*, which encodes intact *ngtA*^41,42^. Accordingly, we speculate that functional NgtA is maintained in environmental lineages due its utility during colonization of a macrophage-like intracellular habitat within eukaryotic microbes. In support of an intracellular environment representing the natural niche of diverse *Francisella* species, species that encode functional NgtA also encode the host cell-targeting type VI secretion system associated with the *Francisella* pathogenicity island^43-45^.

Unlike the GSH uptake mechanisms characterized in other bacterial pathogens, which consist of ABC transporters, intact GSH import in *Francisella* is mediated by an MFS transporter. The consequences of this are unclear; however, one difference between the transporter types is the steepness of the concentration gradient of GSH that each can overcome. ABC transporters rely on ATP and can achieve transport across gradients much steeper than those achievable with MFS transporters, which can only overcome concentration gradients equivalent to those of the coupling ions^46^. Interestingly, the bacteria in which ABC transporters for GSH have been identified, including *S. pneumoniae* and *H. influenzae*, reside in extracellular host-associated niches, where the concentration of GSH is much lower than the intracellular habitat of *F. novicida*^8,9,47^. Thus, differences in the GSH concentration encountered in the different primary habitats these organisms colonize appears to correlate with the GSH uptake mechanism employed, in a manner consistent with the energetics of uptake by each route.

While our data clearly demonstrate that NgtA is capable of transporting GSH and suggests it does not play a role in Cys–Gly import, we have not defined the extent of its physiologically relevant substrates. To our knowledge, the only other MFS protein previously shown to transport GSH is Gex1 of yeast^48^. While Gex1 can export GSH, its primary function appears to be related to cadmium detoxification via the extrusion of GSH-cadmium conjugates. This raises the possibility that NgtA could transport substrates beyond GSH. Several other members of the Pht family of transporters to which NgtA belongs facilitate uptake of amino acids that are limiting during intracellular growth of *Francisella* and *Legionella*^15-17^. Candidate additional substrates for NgtA could include other γ-Glu amide bonded molecules, or a broader range of oligopeptides, such as those transported by members of the proton-dependent oligopeptide transporter class of MFS proteins^49^.

During both growth in GSH medium and intramacrophage replication, we find that either Ggt or NgtA and Cgc1-mediated degradation of GSH are sufficient to support growth of *F. novicida*, raising the question as to why the two pathways are maintained in parallel in many strains. GSH plays several important roles beyond serving as a source of nutrients, including redox buffering, combating oxidative stress and detoxifying metals and xenobiotics^50^. Import of intact GSH via NgtA could thus provide a source of GSH under conditions where *de novo* biosynthesis may provide an insufficient supply of the intact molecule to counteract particular stresses. For the non-pathogenic *Francisella* species in which we find intact *ngtA*, one such condition may be encountered during replication within a protozoan host. These organisms employ many of the same mechanisms for killing phagocytosed bacteria as macrophages, including generation of a reactive oxygen burst^39^. Interestingly, laboratory studies employing model amoeba strains suggest that in these hosts, *Francisella* species replicate within the vacuole where the oxidative burst is delivered, rather than escaping to the cytosol as in mammalian cell infections^51-54^. Additionally, virulent strains of *Francisella* can limit the oxidative burst within cells they infect, by mechanisms that are not yet fully characterized^55-57^; it remains to be determined if these are conserved in other non-pathogenic species. We speculate that NgtA may provide a means of rapidly acquiring GSH for *Francisella* species that must contend with acute episodes of oxidative stress.

If the primary role of NgtA is to enable import of intact GSH for non-nutritional uses, the question then arises as to why *Francisella* species additionally encode an intracellular enzyme for GSH degradation, Cgc1. In eukaryotic cells, constitutive degradation of GSH by intracellular enzymes contributes to GSH homeostasis. In yeast, this is mediated by the Dug complex, which shares the same predicted enzymatic function as Cgc1, whereas in mammalian cells, constitutive turnover of GSH appears to be mediated by the γ-glutamyl cycotransferase enzyme ChaC2^23,58,59^. The ChaC2 homologs that have been characterized to date exhibit a very slow rate of GSH turnover, which has been suggested to be important for preventing unchecked depletion of intracellular GSH levels^59^. We similarly observed a low rate of GSH turnover by purified Cgc1. While this observed rate of turnover may be the result of our *in vitro* assay conditions, it is consistent with Cgc1 playing a role in GSH homeostasis. Of note, *Francisella* spp. also encode a homolog of ChaC2, but this protein localizes to the periplasm. Additionally, strains lacking ChaC exhibit pleitropic phenotypes^11^, and the corresponding gene was not a strong hit in our Tn-seq screen for genes important in GSH utilization, suggesting its role in GSH catabolism is likely a minor part of its overall function in *Francisella*.

A previously missing component of the GSH utilization pathway in *Francisella* is the means by which the tripeptide crosses the outer membrane. Here, we provide evidence that FupA provides this function by acting as a porin. Our findings challenge the prior assertion that FupA serves as a high affinity ferrous iron transporter^32^. Upon reexamination, two pieces of published data support our conclusion that FupA functions as a general porin: *i*) unlike typical high affinity transport mechanisms, FupA expression is not induced by limiting iron, and *ii*) growth of *F. tularensis* Δ*fupA* is reduced in minimal media regardless of the concentration or type of iron supplied^30,32^. Additionally, we note that in other Gram-negative species, porins related to OmpC or OmpF, which are absent in *Francisella* spp., allow passive entry of Fe^2+^ that is then imported across the inner membrane by high affinity transporters^60^. Together with our discovery of the NgtA and Cgc1-mediated pathway for GSH uptake and degradation, our identification of the role of FupA in GSH import allows us to construct a substantially revised model for GSH catabolism in *Fransicella* that highlights the central importance of this molecule for this diverse group of organisms.

## Acknowledgements

The authors wish to thank members of the Mougous laboratory for helpful suggestions. This work was supported by the NIH (R01AI145954 to JDM, SLD, ShJS and JC, P30 DK089507 to StJS), the Defense Advanced Research Projects Agency Biological Technologies Office Program: Harnessing Enzymatic Activity for Lifesaving Remedies (HEALR) under cooperative agreement No. HR0011-21-2-0012 (to JDM and JJW), and the Cystic Fibrosis Foundation (SINGH19R0 to StJS).

## Figure Legends

**Figure S1.**
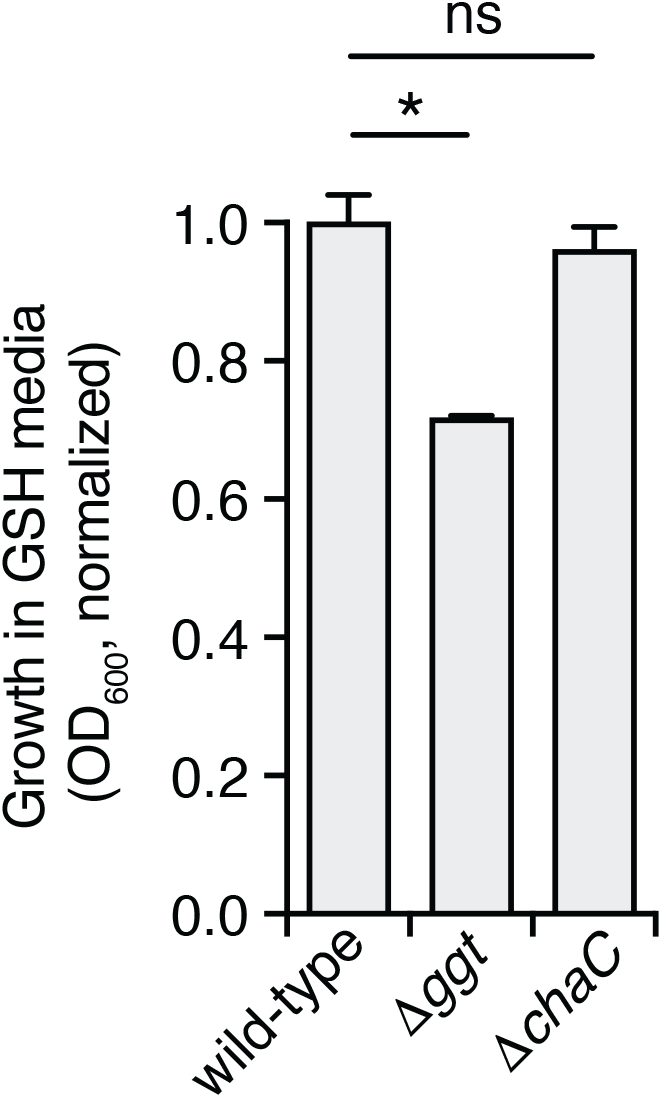
*F. novicida* Δ*ggt* exhibits a growth defect in media containing limiting GSH. Normalized growth yields in GSH medium (20 μM GSH) of the strains. Data are represented as means ± s.d. Asterisks represent statistically significant differences (Unpaired two-tailed student’s t-test.; *p<0.05, ns, not significant). Related to Figure 1.

**Figure S2.**
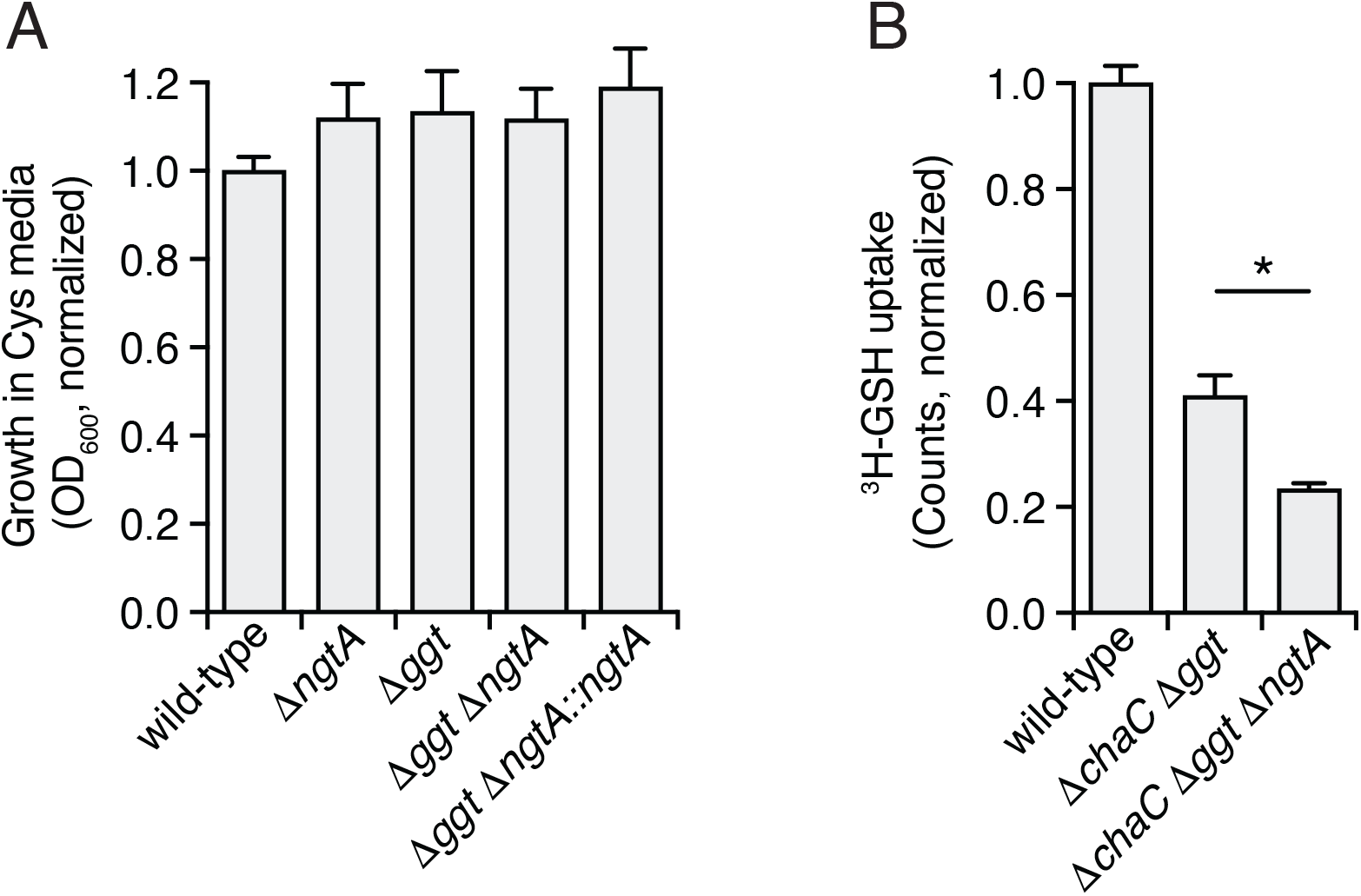
*ngtA* mutants do not display growth defects in cysteine medium and NgtA facilitates GSH import in the absence of periplasmic GSH cleavage. (A) Normalized growth yields of the indicated *F. novicida* strains after 36 hrs in defined medium containing cysteine. (B) Quantification of the level of ^3^H-GSH uptake in the indicated strains of *F. novicida* after 45 min incubation. Data are represented as means ± s.d. Asterisks represent statistically significant differences (Unpaired two-tailed student’s t-test.; *p<0.05, ns, not significant). Related to Figure 2

**Figure S3.**
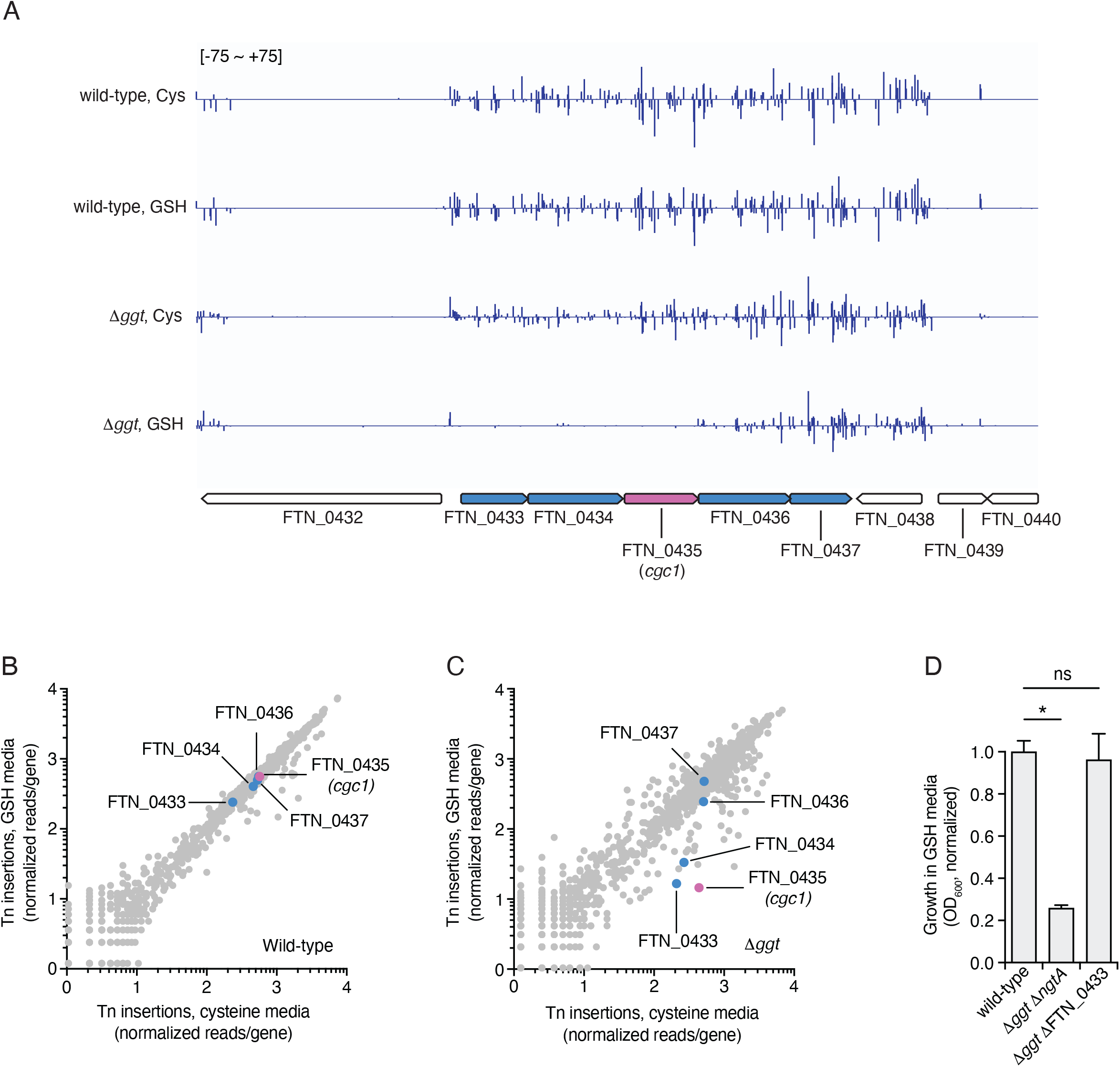
Transposon insertions in FTN_0433 and FTN_0434 cause polar effects on *cgc1*. (A) Transposon insertion profiles for the indicated genetic regions of *F. novicida* (bottom) during growth on media containing GSH or cysteine (cys) as a sole cysteine source. Line height represents the relative abundance of sequencing reads at that position. *cgc1* is highlighted in purple. Other genes in the *cgc1* operon are highlighted in blue. (B,C) Results of the Tn-seq screen shown in Figure 1 reproduced with the genes in the same operon as *cgc1* highlighted. Colors correspond to (A). (D) Normalized growth yields in GSH medium of the indicated strains of *F. novicida*. Data are represented as means ± s.d. Asterisks represent statistically significant differences (Unpaired two-tailed student’s t-test.; *p<0.05, ns, not significant). Related to Figure 3.

**Table S1**. Normalized transposon insertion frequency from a library constructed in wild type *F. novicida* grown in GSH or cysteine media

**Table S2**. Normalized transposon insertion frequency from a library constructed in *F. novicida* Δ*ggt* grown in GSH or cysteine media

## Methods

### Bacterial strains and growth conditions

Bacterial strains used in this study include *Francisella tularensis* subspecies *novicida* U112 (*F. novicida*) and *F. tularensis* subsp. *novicida* MFN245 (*F. novicida* MFN245, both are gifts from Colin Manoil, University of Washington, Seattle, WA), *F. tularensis* subsp. *holarctica* LVS (*F. tularensis* LVS, provided by Karen Elkins, Food and Drug Administration, Rockville, MD), *Escherichia coli* strain DH5α (*E. coli* DH5α, Thermo Fisher Scientific, Cat#18258012), *E. coli* strain BL21 (DE3) (*E. coli* BL21, EMD Millipore, Cat#69450). *F. novicida* strains were routinely grown aerobically at 37°C in tryptic soy broth or agar supplemented with 0.1% (w/v) cysteine (TSBC or TSAC). *F. tularensis* LVS was grown aerobically at 37 °C in either liquid Mueller-Hinton broth (Difco) supplemented with glucose (0.1%), ferric pyrophosphate (0.025%), and Isovitalex (2%) (MHB) or on cystine heart agar (Difco) supplemented with 1% hemoglobin (CHAH). For selection, antibiotics were used at the following concentrations: kanamycin at 5 μg/ml (LVS), 15 μg/ml (U112) or 50 μg/ml (*E. coli*), carbenicillin at 150 μg/ml, and hygromycin at 200 μg/ml. *F. novicida* strains were stored at -80°C in TSBC supplemented with 20% (v/v) glycerol. *E. coli* strains were stored at -80°C in LB supplemented with 15% (v/v) glycerol.

### Strain and plasmid construction

Deletion mutations and in cis gene complementation strains of *F. novicida* were generated via allelic exchange as described previously^61^. Briefly, sequences containing ∼1000 bp flanking the site of deletion or the complementation allele were amplified by PCR and cloned into the BamHI and PstI sites of the vector pEX18-pheS-km using Gibson assembly^44^. Naturally competent *F. novicida* was prepared by back-diluting overnight cultures 1:100 in 2 mL TSBC, growing for 3 hrs at 37°C with shaking, harvesting by centrifugation, and resuspending in 1 mL *Francisella* transformation buffer (per liter; L-arginine, 0.4 g; L-aspartic acid 0.4 g; L-histidine, 0.2 g, DL-methionine, 0.4 g; spermine phosphate, 0.04 g; sodium chloride, 15.8 g; calcium chloride, 2.94 g; tris(hydroxymethyl) aminomethane 6.05 g) ^61^. Approximately 1 μg of pEX18-pheS-km-based deletion or complementation plasmid was added to freshly prepared competent cells. Bacterial suspensions were then incubated at 37°C with shaking for 30 min, followed by addition of 2 mL TSBC and an additional 3 hrs of incubation. Transformants were selected by plating on TSAC with kanamycin. The resulting merodiploids were grown overnight in non-selective TSBC, diluted 1:100 into Chamberlain’s defined medium (CDM)^62^ containing 0.1% p-chlorophenylalanine (w/v) and allowed to grow to stationary phase. Cultures were then streaked onto TSAC, colonies were patched onto TSAC with and without kanamycin to test for kanamycin sensitivity, and kanamycin sensitive colonies were screened for mutations by colony PCR.

The *fupA* complementation strain (Δ*fupA Tn7:Pbfr-fupA*) was constructed using the mini-Tn7 system^63^. Briefly, the *fupA* gene was amplified from *F. novicida* by PCR and cloned using Gibson assembly into the pMP749 plasmid along with the sequence encoding the bacerioferritin promoter (Pbfr) for high constitutive expression^63,64^. Using natural transformation as described above, the resulting plasmid, pMP749:*Pbfr -fupA*, was transformed into plasmid compatible strain *F. novicida* MFN245 carrying the helper plasmid encoding the transposase for Tn7 integration, pMP720. Tn7 integrants were selected on kanamycin, and colonies were screened for the presence of the inserted transposon at the *glmS* locus using PCR. To transfer the *Tn7:Pbfr-fupA* insertion from *F. novicida* MFN245 to *F. novicida* U112, genomic DNA was prepared from *Tn7:Pbfr-fupA* MFN245 strains and 10 ng was used to transform competent U112, prepared as described above. For expressing NgtA^NOV^ in *F. tularensis* LVS, the gene was amplified from *F. novicida* using PCR and cloned into the expression plasmid pF behind the constitutive *groEL* promoter ^65^. Empty pF plasmid or pF-*ngtA*^NOV^ were electroporated into wild-type or Δ*ggt F. tularensis* LVS as previously described^66^, and transformants were selected by plating on CHAH with kanamycin.

For constructing protein expression plasmids pET-28b(+)::6xHis_Cgc1 or pET-28b(+)::6xHis_Cgc1^C97A^, *cgc1* and *cgc1*^*C97A*^ were amplified by PCR and cloned into the NdeI and BamHI sites of the vector pET-28b(+) using Gibson assembly

### Transposon mutant library generation

Transposon mutant libraries containing 100,000 to 300,000 Mariner transposon insertions in *F. novicida* wild-type and Δ*ggt* were constructed using delivery plasmid pKL91^11^. The plasmid was delivered via natural transformation as described above, cells were allowed to recover 2 hrs and then plated on TSAC with kanamycin. Plates were incubated for 20 hrs at 37°C and the resulting kanamycin-resistant colonies were scraped and resuspended in CDM broth lacking a cysteine source (CDM-Cys). Each library was washed 4x times with CDM-Cys broth prior to freezing of aliquots containing ∼10^6^ CFU each in CDM-Cys with 20% (v/v) glycerol at -80°C.

### Tn-seq screen

For each genetic background, two aliquots of the respective transposon libraries were thawed and used as the inocula for 50 mL CDM-Cys media. Cultures were placed at 37°C with shaking for 2 hrs prior to addition of 100 μM cysteine or GSH, followed by incubation for 20 hrs at 37°C with shaking. Cells were then collected via centrifugation and genomic DNA was extracted using the DNeasy Blood and Tissue Kit (Qiagen). Sequencing libraries were generated essentially as described^67^. In brief, 3 μg DNA from each sample was sheared to ∼300 bp on a Covaris LE220 Focused-Ultrasonicator followed by DNA-end repair, terminal C-tailing and of amplification of the transposon-genome junctions by two rounds of PCR. The first round employed the tranposon-specific primer 5’-CATCCTGACGGATGGCCTTTTTGCGTTTCTACC-3’ and olj376, and second round employed the transposon-specific primer5’-AATGATACGGCGACCACCGAGATCTACACTCTTTCGGCATACGAAGACCGGGGACTTATCATCCAACCTG-3’ and distal primers as described^67^ to add unique de-multiplexing barcodes per sample. The libraries were pooled and sequenced using custom sequencing primer 5’-GCATACGAAGACCGGGGACTTATCATCCAACC-3’ and Read1_SEQ primer as described^67^ by single-end 150 bp sequencing with a single index read on an Illumina MiSeq at 11 pM density with 15% PhiX spike-in.

### Tn-seq data processing

Custom scripts^67,68^ (https://github.com/lg9/Tn-seq) were used to process the Illumina sequencing reads and map sites of transposon insertion. First, reads from each sample were filtered for those displaying transposon end sequence (the sequencing primer was designed to anneal five bases from the end of the transposon). The filtered reads were mapped to the genome after removing the transposon end sequence. Reads per unique mapping position and orientation were tallied and read counts per gene were calculated by summing reads from all unique sites within each gene’s ORF except those within the 5’ 5% and 3’ 10% (insertions at gene termini may not be fully inactivating). Gene counts per sample were normalized based on a comparison between all samples of the median reads per gene per gene length for genes with insertions in all samples, as described^68^.

### Bacterial growth assays

To examine the proliferation of *F. novicida* on different cysteine sources, strains were first grown overnight in CDM with 57 μM cysteine at 37°C with shaking. Cells were washed three times and resuspended in CDM lacking cysteine (CDM-cys) followed by cysteine starvation for 2 hrs at 37°C with shaking. Cultures were then diluted to an OD_600_= 0.1 in CDM-Cys supplemented with either GSH, cysteine, or Cys-Gly at 100 μM. Cultures were transferred into a 96-well plate and incubated in a plate reader at 37°C. OD_600_ measurements were taken following a 2 s shake every 10 min. Final growth yields reported represent the OD_600_ obtained after 36 hrs of growth. For *F. tularensis* LVS, strains containing either empty pF plasmid or pF-*ngtA*^NOV^ were first grown overnight in CDM with 1 μM cysteine and then washed and back-diluted to starting OD_600_ = 0.1 in CDM-Cys supplemented with GSH at 100 μM concentration. Cultures were then incubated at 37°C with shaking, and the growth yield was determined by measuring OD_600_ at 16 hrs.

### GSH uptake assays

The indicated strains of *F. novicida* were grown to mid-log phase at 37°C in CDM-Cys liquid medium supplemented with 57 μM cysteine. Cells were spun down, washed three times in uptake buffer (25 mM Tris pH 7.5, 150 mM NaCl, 5 mM glucose), then concentrated 20-fold. The OD_600_ was measured and normalized to OD_600_ = 10. Reactions were established containing 20 μL cells, 5 μL 100 μM GSH, 0.5 μCi 3H-GSH ([Glycine-2-3H]-Glutathione,>97%, 50μCi, PerkinElmer), and 25 μL uptake buffer and incubated for 45 min at 37°C followed by quenching with 1 mL ice-cold uptake buffer. Cells were then pelleted by centrifugation, washed three times in 1 mL cold uptake buffer, then resuspended in 50 μL uptake buffer. Samples were then added to scintillation cocktail (National Diagnostics Ecoscint Ultra) and counts were measured over 1 minute on a scintillation counter (Beckman, LS6500).

### Protein expression and purification

For protein expression, overnight cultures of *E. coli* BL21 carrying pET-28b(+)::6xHis_Cgc1 or pET-28b(+)::6xHis_Cgc1^C97A^ were back diluted 1:500 in 2xYT broth and grown at 37°C until the OD_600_ reached 0.4 ∼ 0.6. Protein expression was then induced by the addition of IPTG (IPTG), and cultures were then incubated with shaking at 18°C for 18 hrs. Following this incubation, cells were collected by centrifugation and resuspended in buffer containing 500 mM NaCl, 50 mM Tris-HCl pH 7.5, 10% glycerol, 5 mM imidazole, 0.5 mg/ml lysosome, 1 mM AEBSF, 10 mM leupeptin, 1 mM pepstatin, 1 mU benzonase, and 5 mM β-mercaptoethanol (BME). Cells were disrupted by sonication and cellular debris was removed by centrifugation at 45,000 x g for 40 min. Lysates were run over a 1 mL HisTrap HP column on an AKTA FPLC purification system to purify the His-tagged proteins. The bound proteins were eluted using a linear imidazole gradient from 5 mM to 500 mM. The purity of each protein sample was assessed by SDS-PAGE and Coomassie brilliant blue staining, and fractions with high purity were concentrated using a 10 kDa cutoff filter. Protein samples were further purified by running over a HiLoad™ 16/600 Superdex™ 200 pg column equilibrated in sizing buffer (300 mM NaCl, 50 mM Tris-HCl pH 7.5, and 1 mM TCEP). Again, the purity of each fraction was assessed by SDS-PAGE and Coomassie brilliant blue staining. Fractions of the highest purity were pooled, concentrated, and utilized in biochemical assays.

### In vitro glutaminase assays

Glutamine amidotransferase activity of purified Cgc1 and Cgc1^C97A^ toward different substrates was assayed in vitro using a glutamate detection kit. 1 μM purified protein was mixed with GSH, glutamine or buffer alone (300 mM NaCl, 50 mM Tris-Cl (pH 8.5)) in 50 μl reactions mixes and incubated 1 hr at at 37 °C. The reactions were stopped by heating at 95 °C for 5 min to inactivate the enzyme. After inactivation, 45 μL of reaction mixtures were added to 100 μL of glutamate detection reaction mix (Abcam Glutamate Assay Kit, ab83389) in a 96-well plate. The reaction mixture was incubated for 5 min at RT followed by 5 min at 37°C. The color change is proportional to the glutamate generated and was measured at A450 in a plate reader. Reported glutamate concentrations were calculated by subtracting background absorbance readings from a no enzyme control.

### H_2_O_2_ sensitivity assays

To monitor H_2_O_2_ tolerance levels, strains of *F. novicida* were grown at 37°C in CDM medium to mid-log (OD_600_= 0.4-0.6). Cultures were diluted to an OD_600_= 0.1 and cell viability was assayed via plating for CFU enumeration. 1.5 mM of H_2_O_2_ was then added and the cultures were placed at 37°C with shaking. After 30 mins and 60 min, samples were collected for CFU enumeration. Survival rates were calculated by comparing the CFU numbers pre and post-H_2_O_2_ exposure.

### Murine bone marrow-derived macrophage generation

Murine bone marrow-derived macrophages (BMMs) were differentiated from bone marrow of female, 6-12 -weeks-old C57BL/6J mice (Jackson Laboratory) for 5 days in non-tissue culture-treated Petri dishes at 37°C under 10% CO_2_ in Dulbecco’s Modified Eagle’s Medium, containing 1g/L glucose, L-glutamine and sodium pyruvate (DMEM) supplemented with 10 % fetal bovine serum (FBS) and 20% L-929 mouse-fibroblast conditioned medium (L-CSF). 5 days post-plating, non-adherent cells were washed out with ice-cold phosphate buffered saline (PBS) and the differentiated BMMS were incubated for 10 min in ice-cold cation-free PBS (Corning) supplemented with 1 g/L glucose, detached by pipetting and harvested by centrifugation for 7 min at 200xg/25°C. Pelleted cells were resuspended in BMM complete medium (DMEM, 10% FBS, 10% L-CSF) and plated at a density of 5×10^4^ cells/well in 24-well, tissue culture-treated plates followed by incubation for 48 h at 37°C under 10% CO_2_ with replenishment of BMM complete medium at 24 hrs post-plating.

### Macrophage infection assays

48 hrs post-plating, BMMs were infected with mid-log phase *F. novicida* at a multiplicity of infection (MOI) of 1 in pre-chilled BMM complete medium. Bacterial uptake was synchronized by centrifugation for 10 min at 400xg/4°C after which the plates were immediately placed in a water tray pre-warmed to 37°C and incubated for 30 min at 37°C under 10% CO_2_. Following incubation, extracellular bacteria were removed by 4 washes with plain DMEM medium pre-warmed to 37°C, the complete BMM medium was replenished, and the plates were placed back at 37°C under 10% CO_2_. At 2 hrs and 24 hrs post-infection (p.i.) the BMMs were rinsed 3 times with sterile PBS and lysed in PBS/0.1% sodium deoxycholate (Sigma), followed by serial dilution in sterile PBS and plating on TSAC plates for CFU enumeration. Bacterial growth at 24 h p.i. was normalized to the CFU counts obtained at 2 h p.i.

### Mice

C57BL/6J mice used in this study were purchased from Jackson Labs. Mice were maintained under SPF conditions ensured through the Rodent Health Monitoring Program overseen by the Department of Comparative Medicine at the University of Washington. All experiments involving mice were performed in compliance with guidelines set by the American Association for Laboratory Animal Science (AALAS) and were approved by the Institutional Animal Care and Use Committee (IACUC) at the University of Washington.

### *F. novicida* inoculum preparation and intranasal infection

*F. novicida* inoculum was prepared as described previously^61^. Briefly, 3 mL TSBC was inoculated with each *F. novicida* strain and incubated aerobically for 18 h at 37°C with shaking. After overnight growth, cultures were adjusted to OD_600_ =1 in TSBC, diluted 1:1 with 40% glycerol in TSBC (20% v/v final glycerol concentration), aliquoted and stored at -80°C. The post-freeze titer of each stock was determined by culturing on TSAC. Just prior to infection, an aliquot of each strain was quickly thawed at 37°C and diluted in sterile 1X PBS to ∼100 CFU in 30 μL (∼3.3 × 10^3^ CFU/mL).

Mice were infected with indicated *F. novicida* strains by intranasal instillation (30 μL total) under light isoflurane anesthesia. Mice were weighed just prior to and 48 hrs post infection. After 48 hrs mice were euthanized with CO_2_. The lungs and spleens were harvested in 5 mL lysis buffer (0.1% IGEPAL, in 1x PBS sterile filtered) and homogenized using a Tissue Teaeror Homogenizer (BioSpec Products). Organ homogenate was serially diluted into 1X PBS and dilutions plated on TSAC. Plates were incubated at 37°C overnight aerobically. Colonies were counted and CFU per organ calculated.

### NgtA sequence and phylogenetic analysis

To generate a phylogeny, NgtA homologs were identified by collecting the top 5,000 hits from Psi-BLAST, then aligned using the Clustal Omega plug in of Geneious Prime (Dotmatics). Positions with gaps present in at least 30% of sequences were masked in the alignment, and then a neighbor-joining phylogeny was constructed using the Geneious Tree Builder. This phylogeny included both Pht family members and a number of clades of related MFS family proteins from other subfamilies. Non-Pht family clades were eliminated by performing additional BLASTp searches with representatives from each clade that did not contain a previously characterized Pht family member; clades were eliminated when these sequences had higher percent identity matches with other MFS transporter families than with the closest Pht family member. This yielded a set of 1,043 sequences that were re-aligned and masked as described above, and used to construct a new neighbor joining phylogeny.

Inactivating mutations in *ngtA* coding sequences were identified by performing a tBLASTn search with NgtA, limited to the Thiotrichales. All protein sequences obtained were filtered to remove those sharing <50% identity with NgtA of *F. novicida*, as these were found to represent other Pht family members. Remaining protein sequences were then aligned. In cases where a premature stop codon had been introduced into the coding sequence of *ngtA*, our tBLASTn search retrieved multiple hits, which manifested as truncated sequences in the protein sequence alignment. For *F. endociliophora*, the absence of *ngtA* was confirmed by performing a BLASTp search with NgtA against the complete genome, and by examining the conserved genomic location where *ngtA* is encoded in other strains.

